# Imputing the parental origin of the sibling’s haplotype from parental phenotypes

**DOI:** 10.1101/2022.07.09.499429

**Authors:** Yanyu Liang

## Abstract

To recruit cases for late-onset disease study is challenging since these diseases occur in elder people. Moreover, typically we have a very limited number of late-onset disease cases in Biobank data. But, on the other hand, the parental disease status may be available by questionnaire. Because of this, methods have been developed to utilize parental disease status instead Liu et al. (2017); Hujoel et al. (2020). In these approaches, the late-onset phenotype of the participant is imputed from parental statuses. And, downstream, a genome-wide association study (GWAS) is performed using the participant’s genotype and imputed phenotype. In this paper, we take another view on utilizing parental phenotypes. We treat this problem as missing parental genotype rather than missing participant’s phenotype. First, we propose an imputation scheme to infer the parental origin of the participant’s genotype from a collection of extra parental phenotypes (non-focal phenotypes) and the participant’s genotype. Second, we propose a computationally efficient approach to incorporate the imputed parental origin information into the downstream GWAS. We explore the feasibility of the proposed two-step approach on simulated and real data. And we derive the power increase of GWAS as a function of imputation quality. These results indicate that the imputation scheme needs about 100 non-focal phenotypes to achieve enough accuracy to facilitate the GWAS downstream.

## 1 Introduction

Late-onset diseases, by definition, occur in elder people, which increases the difficulty to recruit cases for GWAS. Nowadays, biobank data has become an important data source for GWAS because of its ability to collect genome and phenome of a large population. And the GWASs of some complex diseases with relative high prevalence in a general population have achieved great successes in utilizing biobank-scale data set. However, the biobank usually has limited number of the late-onset disease cases due to the fact that the participants are typically at the middle-age and healthy. For instance, looking at late-onset Alzheimer’s disease in UK Biobank. There are about 20% of UK Biobank participants older than 65 at the age of recruitment Sudlow et al. (2015) and around hundreds of late-onset Alzheimer’s disease cases. The reason of short of cases is two-fold. First, there are only about 100,000 elder people to start with without specifically targeting Alzheimer’s disease research. Second, the recruitment could be biased towards healthier participants, which may give rise to a even lower prevalence than the Alzheimer’s diease prevalence in a general population Isik (2010).

On the other hand, the parents of participants usually were or have been at the age of high prevalence of late-onset disease. It means that the parents of participants are more ideal objects to work with since the disease of interest is more prevalence. And when looking at the participant’s parents, we don’t need to limit the scope to elder participants. Moreover, in principle, the parental case/control status of the late-onset disease is possibly achievable from the participants via questionnaire. For example, UK Biobank has collected 12 parental phenotypes via web-based questionnaires, and, in particular, there are over 26,000 paternal Alzheimer’s disease cases and over 49,000 maternal Alzheimer’s disease cases Sudlow et al. (2015). In comparison, a recent GWAS which is dedicated to late-onset Alzheimer’s disease Kunkle et al. (2019) have around 35,000 cases.

Motivated by the potential richness of late-onset disease cases in parental phenotypes, methods have been developed to leverage parental phenotypes for the corresponding GWAS. In this context, the challenge is that we only have access to the genotype of the child (the participant) and the phenotype of the parents. The existing approaches treat it as a missing phenotype problem, in which they first impute the missing phenotype. By missing phenotype, it means that the phenotype of the child is not yet observable since the participant has not reached the typical age of onset. And the subsequent GWAS is performed using child’s genotype and imputed phenotype. To impute the child’s phenotype from the parental phenotype, GWAX was proposed to use the parental phenotype as a proxy of the child’s phenotype Liu et al. (2017). More recently, Hujoel et al. (2020) imputed the genetic liability of the child from the family disease history by modeling case/control phenotype with the liability threshold model.

In this paper, we want to take another view of the problem. Here, we treat it as a missing genotype problem instead. Specifically, we consider parent’s genotype as missing and we want to, first, impute parent’s genotype and, second, perform GWAS with observed parental phenotype along with the imputed genotype of the parents. Since one of the child’s alleles is from father and the other is from mother, essentially, the imputation is about inferring which allele is from father and which allele is from mother. Moreover, considering that we typically have access to phased genotype, the imputation problem could be further reduced to inferring the parental origin of the two haplotypes of the child. To infer the haplotype origin, we need to leverage the information buried in the parental phenotypes other than the one of interest (we call them non-focal phenotypes below). In particular, each haplotype carries the genetic risks of these non-focal phenotypes, which could provide information on which haplotype is more likely to come from which parent. Intuitively, if the genetic risks carried by one haplotype resemble the corresponding observed maternal phenotypes, then, this haplotype is more likely to come from mother as compared to father. A concrete example of the intuition is shown in Figure 1.

**Figure 1:**
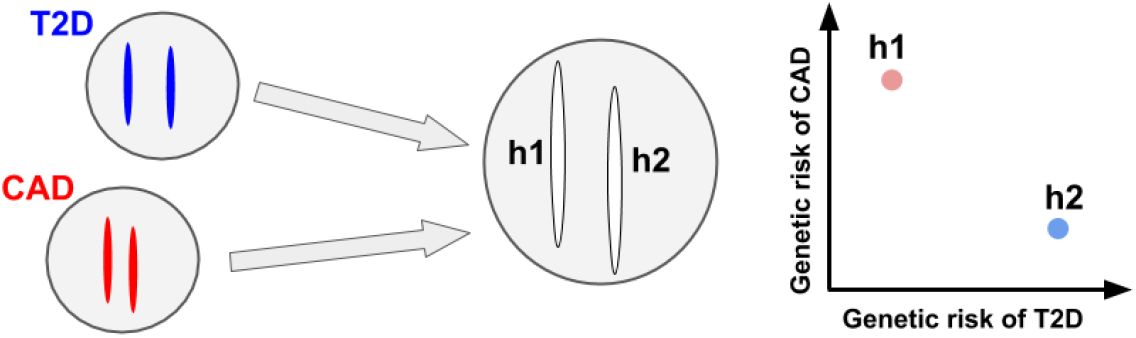
An example on imputing haplotype origin from non-focal phenotypes of parents and genetic risks carried in child’s haplotypes. In this figure, a circle represents a cell with two chromosomes inside representing the sister chromosomes of a diploid (in practice, there are 23 pairs and here we only show one chromosome for the sake of simplicity). On the left, the two circles represent the cell from the father and the mother respectively with the chromosomes in father colored in blue and the ones in mother colored in red. And suppose the father is observed to have type II diabetes (T2D) and the mother is observed to have cardiovascular disease (CAD). The circle in the middle represents the cell of the child (the participant in the biobank) where the two chromosomes are labelled as “h1” and “h2”. These chromosomes are uncolored since the haplotype origin is unknown and is the goal of the imputation. On the right, the plot shows the genetic risk of T2D and CAD carried in “h1” and “h2”. Since “h1” carries higher CAD risk and lower T2D risk, which resembles the paternal phenotype, “h1” is colored with light red indicating that it is more likely that “h1” is from the mother. For similar reason, “h2” is colored with light red.

Subsequent to the imputation of haplotype origin, we propose an approach to integrate the imputed haplotype origin into the GWAS of the focal phenotype, which uses parental phenotypes and child’s haplotypes. Our proposed approach includes GWAX approach as a special case at which the two child’s haplotypes are both equally likely to come from each of the two parents. When the quality of the imputation is good, the proposed approach has higher power to identify GWAS signals than GWAX.

In the following sections, we first describe the imputation scheme for the inference of haplotype origin (Section 2.1). Secondly, we describe the proposed GWAS approach that integrates imputation results and give the theoretical results on how the imputation quality affects the power of the downstream GWAS (Section 2.2). And then, we perform simulation to examine the imputation quality at different parameter settings, such as the number of non-focal phenotypes and the heritability (Section 3.1). Similarly, on the basis of simulated data, we test the proposed GWAS approach and verify the theoretical results (3.2). Following the analysis on simulated data, we apply the imputation scheme to trios in the Framingham transcriptome data and perform downsampling analysis to examine the imputation quality on real data (Section 3.3). Finally, we summarize the observations and the insights obtained from the theoretical results along with the applications to simulated and real data, and discuss the potential pitfall and future direction (Section 4)

## 2 Methods

### 2.1 Imputing haplotype origin

#### Modeling setups

We consider the following problem setup. All the variables below are at individual-level and, for simplicity, we omit the indexing on individual. The phased genotype of the child, *H*^1^ and *H*^2^ representing the two haplotypes, are observed. And we observe parental phenotypes, *y^π^* (*π* = father or mother), which is a vector of length *P* (*P* phenotypes in total). Let *Z* be an indicator variable representing the event that haplotype 1 is from father. The goal is to infer *Z* from data.

Let “half-genotype” be half of the genotype where one of the two alleles is included for each variant, which is a weaker definition than the haplotype so that the alleles are not required to be on the same chromosome. To relate the observed phenotype with the child’s haplotypes, we introduce 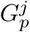 representing the genetic effect of *j*th half-genotype on *p*th phenotype for a given individual. And furthermore, we assume a genetic model as follow.

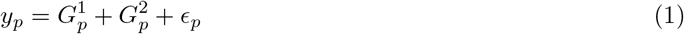

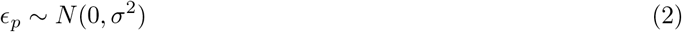

In the context of modeling parental phenotype, from Mendel’s laws, we observe one of the two half-genotypes from a parent, which gives rise to one of the two child’s haplotypes. So, Eq 1 is reduced to

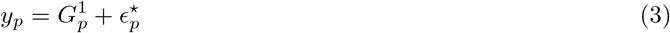

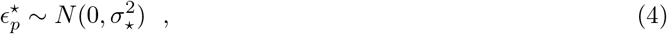

where we assume 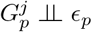 so that 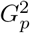 is absorbed into error term. Here the order of the two half-genotypes are arbitrary and we set the one being transmitted to the child as half-genotype 1.

In practice, we don’t observe 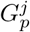 so we further assume that

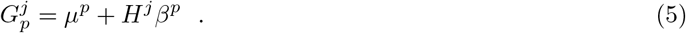

Under this model, we need to fit genetic effect *β^p^* along the way of inference so we refer to this approach as on-the-fly approach. And, moreover, if we have a genetic predictor (obtained from external data), *e.g*. a polygenic risk score model, we can model 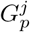 as

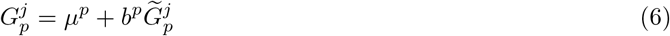

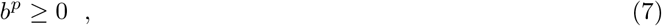

where 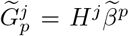 is the predicted value of pth phenotype and *μ^p^*, *b^p^* are scalars which are introduced to account for the different scaling of the phenotype between the observed phenotype here and the one for predictor training. Besides, *b^p^* is set to non-negative to impose the constraint that predictor should at least predict the direction correctly. And we refer to this approach as PRS-based approach.

#### Constructing the likelihood and the imputation

On the basis of Eq 3, Eq 5, and Eq 6, we propose the following likelihood function for the observed data *y*^father^, *y*^mother^, *H*^1^, *H*^2^

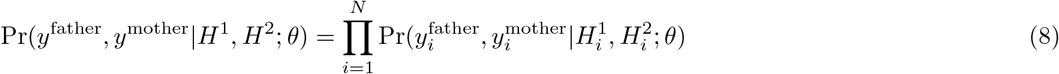

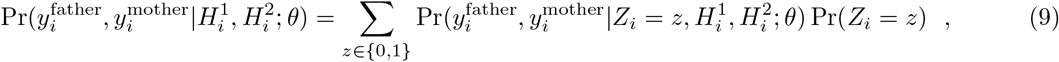

where Pr(*Z_i_* = 0) = Pr(*Z_i_* = 1) = 0.5 for all individuals since the two child’s haplotypes are equally likely to come from father and mother before observing the data. And *θ* represents the model parameter in the genetic models proposed in Eq 5 or Eq 6. Moreover, given *Z_i_* = 1, 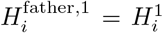 and 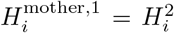. And when *Z_i_* = 0, similar equations follow but *H*^1^ and *H*^2^ flip the order. So, 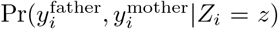 is straightforward to compute.

The goal is to update the distribution of *Z* after observing the data, i.e. to obtain 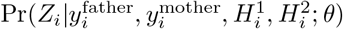. Since the true *θ* is unknown, we propose to plugin the maximum likelihood estimator (MLE) of *θ* under Eq 9. Algorithmically, we use EM algorithm to obtain the MLE where we iterate over updating posterior of *Z* given current parameter *θ*^(*t*)^ and updating *θ* on the basis of the posterior of *Z* (Supplementary Notes 2.1).

In practice, the imputation described above is performed one chromosome at a time. So, the genetic model only carries the heritability of the corresponding chromosome. As there are 22 autosomes with various length and gene density, the per-chromosome heritability is only a fraction of the chip heritability, which also depends on genetic architecture. Because of this, the per-chromosome imputation is less effective than the imputation using all chromosomes jointly. But when fitting all chromosomes jointly, *Z* becomes a 22-length vector and there are 2^22^ possible configurations of *Z*. To resolve this computation burden, we propose an alternative approach which updates one chromosome at a time (Supplementary Notes 2.2).

### 2.2 Integrating imputation results to GWAS

#### Problem overview

Here we consider performing GWAS using parental phenotype and genotype. As we only observe the genotype of the child and the parental genotype is missing, we, at most, observe one of the two alleles for each locus of the parent. So, similar to Eq 3, we consider using allelic test for GWAS Lee et al. (2013). Specifically, we consider modeling *y^π^*|*H*^*π*,1^, *π* = father or mother, via: 1) linear model if quantitative phenotype, and 2) logistic model if case-control phenotype. And *H*^*π*,1^ represents the parental half-genotype that has been transmitted to the child for a given locus.

#### GWAS with softly assigned haplotype origin (soft-GWAS)

In practice, we don’t observe *H*^*π*,1^ directly but we know that *H*^*π*,1^ is either *H*^1^ or *H*^2^. And from the imputation of haplotype origin (Section 2.1), we have some information about which haplotype is more likely to come from which parents. One way to incorporate this piece of information into the model is by modeling Pr(*Y^π^*|*H*^1^, *H*^2^) instead, which, similar to Eq 9, gives rise to the following model

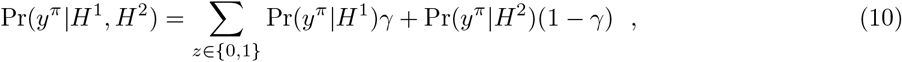

where *γ* is the posterior *Z* obtained from the imputation. To test whether the genetic effect is non-zero, we can use likelihood ratio test to obtain the statistical significance, which corresponds to running an EM algorithm similar to Supplementary Notes 2.1 (see Supplementary Notes 2.3). This approach may result in big computation burden when performing genome-wide analysis.

#### GWAS with imputed half-genotype (imputed-GWAS)

Alternatively, we propose to plugin the posterior mean of the parental half-genotype instead. For instance, in the case of analyzing a quantitative trait using paternal phenotype, we perform linear regression with observed phenotype *y*^father^ and imputed half-genotype 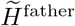 where 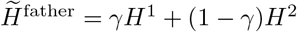. By doing so, we avoid running EM algorithm and the computation complexity is the same as a convention GWAS. And GWAX could be considered as a special case of imputed-GWAS where *γ* = 0.5.

#### The power and bias of imputed-GWAS

Here we focus on the power and bias of the imputed-GWAS approach. For the sake of simplicity, we analyze a simple scenario where phenotype is quantitative. Let *y* be the observed phenotype and 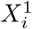 be the true half-genotype. The imputed half-genotype takes the form 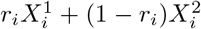. Note that *r_i_* is similar to *γ_i_* but *r_i_* denotes the probability of assigning the haplotype correctly. So, *r_i_* = *γ_i_* if haplotype 1 is from father and *r_i_* = 1 – *γ_i_* otherwise. We assume the genetic model is

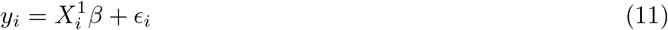

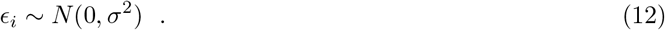

And we rely on linear regression to obtain statistical significance, *i.e*. 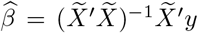. Note that the optimal test is achieved when *r_i_* = 1 for all individuals. And let *T*\* be the test statistic 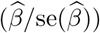 obtained at the optimal. We derive the following results on the bias and power of the imputed-GWAS approach (Supplementary Notes 2.4).

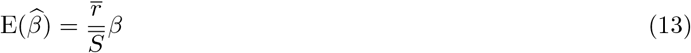

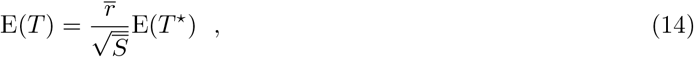

where 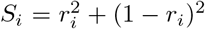 and 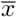 means taking sample mean of variable *x*. As *r* is a measure of imputation quality (the closer to 1, the higher quality), the bias and power both depend on the imputation quality. And the power monotonously increases as *r* increases.

### 2.3 Simulation study of the imputation scheme

#### Simulation to comparing different genetic model and other parameter settings

The goal of this simulation study of the imputation scheme is to see how the imputation quality is affected by heritability, the number of non-focal phenotypes, the choice of the genetic model, and etc. We simulate parental phenotypes, genotypes, and child’s genotype by the following procedure.

1. Simulate parental half-genotypes, *H*^*π*,1^ and *H*^*π*,2^, for *π* = father and mother of each individual. Variants are independently sampled from Bernoulli distribution with minor allele frequency sampled from a uniform distribution*U*[0.05, 0.45].
2. Simulate effect size with *β* ~ (1 – *π*_0_)*δ*_0_ + *π*_0_*N*(0,1) where *π*_0_ = 0.5.
3. Calculate parental genetic effect as *G^π,j^* = *H^π,j^β*.
4. Simulate environmental effect, *ϵ^π^* for each parent where 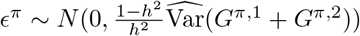.
5. Calculate observed parental phenotype as *y^π^* = *G*^*π*,1^ + *G*^*π*,2^ + *ϵ^π^*.
6. Transmit *H*^father,1^ and *H*^mother,1^ to child with *H*^1^ = *H*^father,1^ and *H*^2^ = *H*^mother,1^.

To simplify the message, we fix the number of variants to 50 and the sample size for imputation to 1,000. Step 2-5 is repeated for 100 parental phenotypes with heritability *h*^2^ = 0.01, 0.05, 0.1, 0.25, 0.5.

To test the PRS-based approach, we also simulate a separate cohort, with 20,000 individuals, for PRS training. To see how PRS quality affects the imputation, we also train PRS using a subset of these individuals (sample size = 5,000 and 10,000).

#### Simulation to examine the PRS-based approach

Here, we are specifically interested in the utility of PRS-based approach in the context that parental transcriptome is available. In this setting, we focus on building the predictor of cis-regulation. Typically, we have hundreds of samples in a transcriptome study and the cis-window contains thousands of variants. So, PRS-approach serves better in this scenario.

To examine the power of PRS-approach in this specific setting, we simulate data by the following procedure.

1. Simulate parental genetically determined gene expression *G*^*π*, 1^ and *G*^*π*,2^.
2. Simulate the parameter *b* in Eq 6 from: 1) *b* = −0.1, 2) *b* = 0.1, 3) *b* ~ *N*(0,1), and 4) *b* ~ max(0, *N*(0,1)).
3. Simulate some covariates *C_m_* ~ *N*(0,1) with effect size *a_m_* ~ *N*(0,1).
4. Calculate *y^π^* = (*G*^*π*,1^ + *G*^*π*,2^)*b*+∑_*m*_ *a_m_C_m_*+*ϵ^π^* where the variance of *ϵ^π^*, 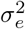, is set such that heritability 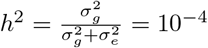,10^-3^, 0.01, 0.05 (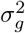 is the genetic variation).
5. For the child, the genetic effect of haplotype 1, *G*^1^, is set to *G*^father,1^ and, similarly, *G*^2^ is set to *G*^mother,1^

For simplicity, we fix sample size to 300, the number of phenotypes to 500, and the number of covariates to 4.

### 2.4 Simulation study of the proposed GWAS approaches

Here we simulate data to test the performance of the proposed GWAS approaches, soft-GWAS and imputed-GWAS. The simulation procedure is as follow which generates one phenotype-genotype pair at a time (individual index is ignored).

1. Simulate *H*^1^ and *H*^2^ from Bernoulli distribution where minor allele frequency is samples from *U* ~ *U*[0.05, 0.45].
2. Simulate phenotype 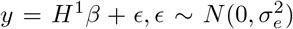 where *β* is either 0 (the null) or 1 (the alternative). And when *β* =1, 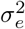 is set such that per-SNP heritability is 0.001; when *β* = 0, 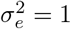.
3. Simulate *γ* ~ *g*(·) where *g*(·) takes the following forms: *δ*_1_, *δ*_0.9_, *δ*_0.1_, *δ*_0.5_, Beta(5, 2), Beta(2, 5), and Beta(2, 2).

We fix sample size to 5,000 and we repeat the procedure 500 times which results in 500 independent phenotype-genotype pairs for the association test. We refer to *δ*. as “constant” distribution and Beta(·, ·) as “beta” distribution. In this simulation, *H*^1^ is the affecting haplotype, so the “optimal” test is obtained at *γ_i_* = 1 which corresponds to *δ*_1_. Moreover, for each distribution type, we include three distributions which have *γ* centered around high values (referred as “high”), around 0.5 (referred as “middle”), and around low values (referred as “low”) So, *δ*_0.9_ and Beta(5, 2) lie in category “high”; *δ*_0.5_ and Beta(2, 2) lie in category “middle”; and *δ*_0.1_ and Beta(2, 5) lie in category “low”.

### 2.5 Analyzing Framingham Heart Study

We obtained genotype and transcriptome data (Joehanes et al. (2017); Zhang et al. (2014)) via dbGap accession number phs000007.v29.p1. The genotype data and gene-level expression quantification have been cleaned-up and processed previously in Wheeler et al. (2016). In brief, the genotype data was pre-phased locally by SHAPEIT Delaneau et al. (2012) and imputed to HRC v1.1 McCarthy et al. (2016) using Michigan Imputation Server Das et al. (2016). Note that under this phasing procedure, the first haplotype of the child is from father and we also verified this result by checking the genetic relatedness between child’s half-genotype and father/mother genotype. And the gene expression data was pre-processed by Affymetrix power tools suite.

We extracted the individuals in the study of whom both the genotype and the trancriptome data were available. Overall 4,838 individuals were selected. We constructed expression matrix with only these individuals being included and then we quantile normalized the expression within each gene. The expression matrix was used to run PEER factor analysis Stegle et al. (2012) to obtain the hidden confounding factors of the experiments where we set the number of factors to 40. We also ran PCA using the genotypes of these individuals via GCTA tools Yang et al. (2011) where the first 20 PCs were kept. These 20 PCs and 40 PEER factors were used as the covariates in the haplotype imputation.

Based on the pedigree information, we extracted the trios where both parents have expression data available and the whole trio have genotype data available. If two trios share the same father and/or mother, we kept only one of them for the analysis. In the end, we collected 266 trios. We applied the PRS-based imputation scheme to these trios where the predicted expression was obtained from elastic net and DAPG weighted elastic net models trained on GTEx V8 whole blood European samples Barbeira et al. (2020a,b). In particular, we ran the imputation for each chromosome, And to examine the power of the imputation, within each chromosome, we downsampled the genes to a fraction of the original number and re-ran the imputation to see how the imputation quality depends on the downsampling fraction.

We defined the genetic relatedness between half-genotype and genotype or genotype and genotype in a unified way. First, we standardized haplotype. For an diploid, we defined 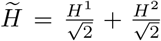 where *H^j^* is the *j*th haplotype. And for a haploid, we defined 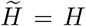. The genetic relatedness is defined as 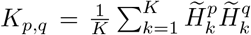 where *p* and *q* represents either diploids (from father’s and mother’s genotype) or haploids (from child’s haplotype). If both *p* and *q* are diploid, this definition is consistent with conventional definition, i.e. *E*(*K_p,q_*) = 1 if *p* = *q* and *E*(*K_p,q_*) = 0 if *p* and *q* are completely unrelated. For the genetic relatedness between haploid 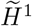 and diploid 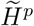,

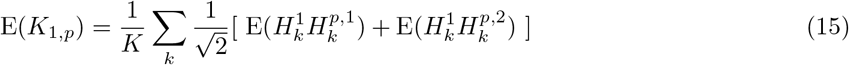

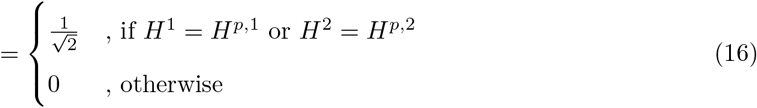

## 3 Results

### 3.1 Examining imputation quality on simulated data

As described in Section 2.1, we proposed two genetic models (Eq 5 and 6) to model parental phenotype *y^π^* given parental half-genotype *H*^*π*,1^. To evaluate these approaches, we examined the imputation performance of these approaches on simulated data where *Z_i_* was set to 1 for all individuals (Section 2.3). We also included an “ideal” approach in the comparison which is, in principle, the “best” imputation we could get under the current model assumption. In the “ideal” approach, we pretend that both the genetic effect and the variance of the environmental effect are observed, which means that *θ* is known. So, the posterior of *Z* can be evaluated under the true parameters, which bypass the uncertainty introduced by fitting model parameter via EM iteration.

We simulated data under different heritabilities and the number of phenotypes included in the simulation also varied. In Figure 2, we show the imputation accuracy (in terms of the probability that the posterior of *Z* is correct) of the three approaches. As the number of phenotypes increases, the imputation accuracy increases for all the three approaches. Similarly, as phenotypes used for imputation are more heritable, *i.e*. the genetic effect captures more observed phenotypic variation, the imputation is more accurate. As expected, the “ideal” approach performs better than the other approaches in all settings. But, across all settings, there are some individuals with accuracy close to zero, especially when the number of phenotype is greater than 10 and heritability is between 0.01 and 0.25. These low-accuracy imputations also appear in the “ideal” approach which suggests that it is due to the intrinsic uncertainty buried in the data.

**Figure 2:**
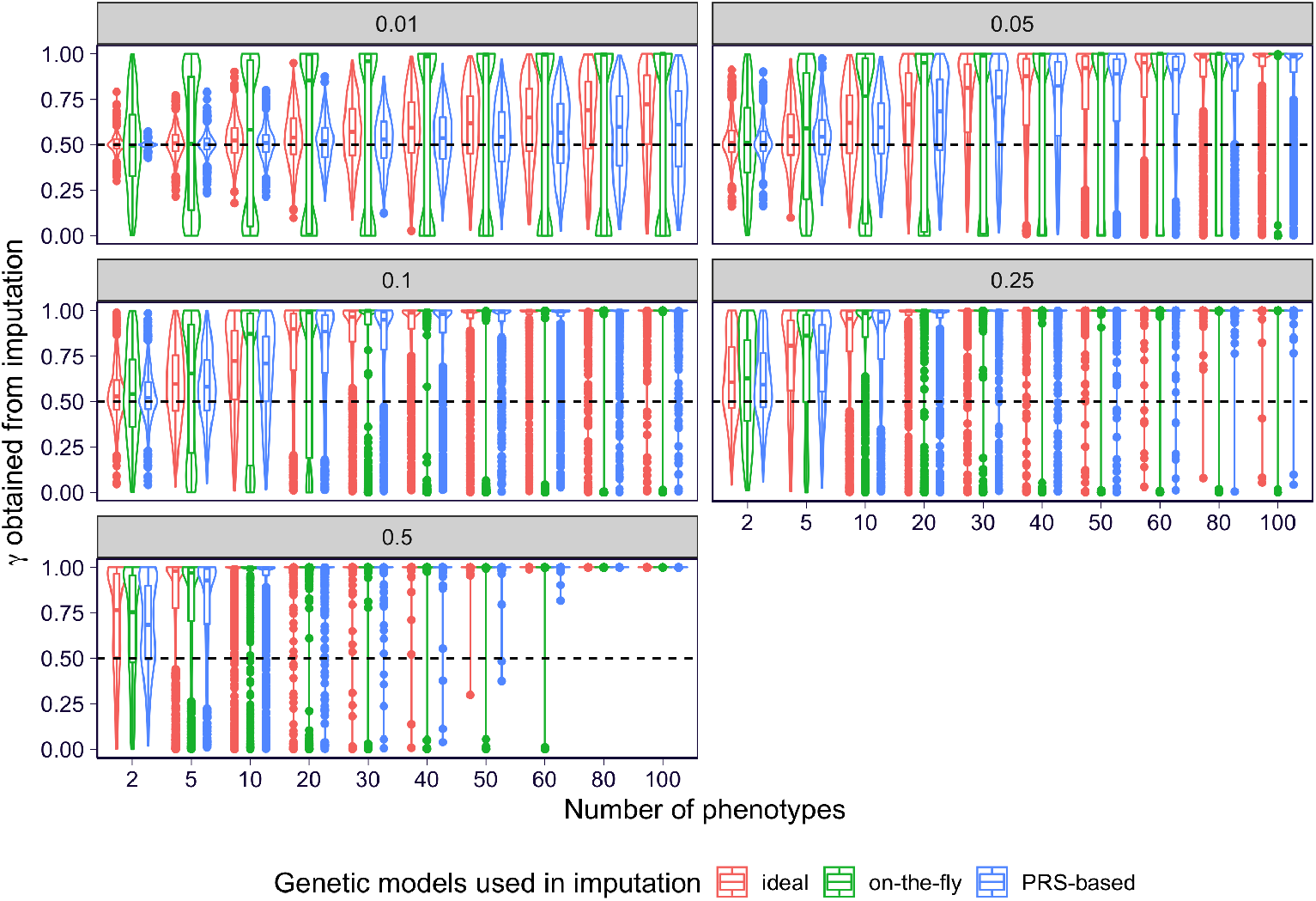
The imputation performance on the basis of different genetic models. The imputation performance under a different heritability is shown in each panel. Within each panel, the number of phenotypes included in the imputation is shown on x-axis and the imputation accuracy (the probability that the posterior *Z* is correctly assigned) is shown on the y-axis. The violin/boxplot contains the results on all of the 1,000 individuals included in the imputation. The results of the “ideal”, on-the-fly, and PRS-based (PRS trained with sample size 10,000) approaches are colored in red, blue, and green respectively.

Comparing the “ideal” approach and the PRS-based approach, they have similar performance with PRS-based approach performs slightly worse which could be attributed to the noise introduced by PRS training. However, the on-the-fly approach performs differently, especially when the heritability is low (0.01 to 0.1). In particular, the on-the-fly approach has close to zero accuracy on many samples, which is most sever when the number of phenotypes is low or the heritability is low. Such bad performance could be attributed to the overfitting issue since the on-the-fly approach has the most flexible genetic model among the three. But, notice that, when the phenotype is highly heritable (stronger signal in the data) or there are many phenotypes being included (the genetic model is more constrained) in the imputation, the on-the-fly approach starts to perform similarly to the other two. And, moreover, the performance of the on-the-fly approach is even better than the PRS-approach when the phenotype is highly heritable. This is due to two reasons. First, the performance of the PRS-based approach is limited by the external PRS model and when there are more samples used in PRS training, the PRS-based approach performs better (Supplementary Figure 1). And, secondly, the on-the-fly approach may learn a better genetic model via the EM iteration.

Then, we performed a another simulation to examine the utility of introducing non-negative coefficient in the PRS-based approach (Eq 6). Here we simulated data under a specific application context where the parental phenotypes are transcriptome profile (Section 2.3). The non-negative constraint on the coefficient b can reduce the amount of error when the true b is non-negative but not so when the b has random or negative sign (Supplementary Figure 2). In practice, if the PRS is at least able to predict the sign to some degree, the non-negative constraint is helpful in the sense of reducing the amount of overfitting.

### 3.2 Verifying the proposed GWAS approach on simulated data

As described in Section 2.2, we proposed two approaches to carry out association study between phenotype and genotype with the imputation results, posterior of *Z* (also called *γ*), being integrated. One of the approach, soft-GWAS, estimates the effect size via EM iteration where the haplotype imputation result is treated as a prior which corresponds to the hidden variable in the EM. And the statistical significance of this approach is obtained from likelihood ratio test. In the other approach, imputed-GWAS, we run association test between the posterior haplotype and the phenotype.

To examine the performance of the two approaches, we simulate some data and run these methods with various kinds of distribution on *γ* which correspond to various imputation qualities (Section 2.4). We included two types of distributions for *γ* where *γ* is either a constant across individuals or drawn from a Beta distribution (labeled as “constant” and “beta”). Moreover, to consider different imputation qualities, we sampled *γ* under three scenarios: 1) *γ* is at low accuracy, *i.e*. posterior *Z* has less than 50% chance to be correct (labeled as “low”); 2) *γ* is at medium accuracy, *i.e*. around 50% chance to be correct (labeled as “medium”); 3) *γ* is at high accuracy, *i.e*. more than 50% chance to be correct (labeled as “high”). We also included the “optimal” scenario where *γ* is 100% accurate for all individuals.

Under the null data, *i.e*. no genetic effect, both soft-GWAS and imputed-GWAS have calibrated p-value (Supplementary Figure 3). When including the genetic effect, the imputed-GWAS and soft-GWAS have similar test statistic *T* (defined as 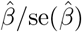) across all settings (Figure 3A). And the similar trends are observed in the estimate effect size 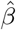 and its standard error as well (Supplementary Figure 4). These results suggest that imputed-GWAS and soft-GWAS performs quite similarly though imputed-GWAS is less interpretable from the first principle but computationally simpler. So, we can reliably treat imputed-GWAS as a good alternative approach to soft-GWAS, which approximates soft-GWAS in a much more computationally feasible manner.

**Figure 3:**
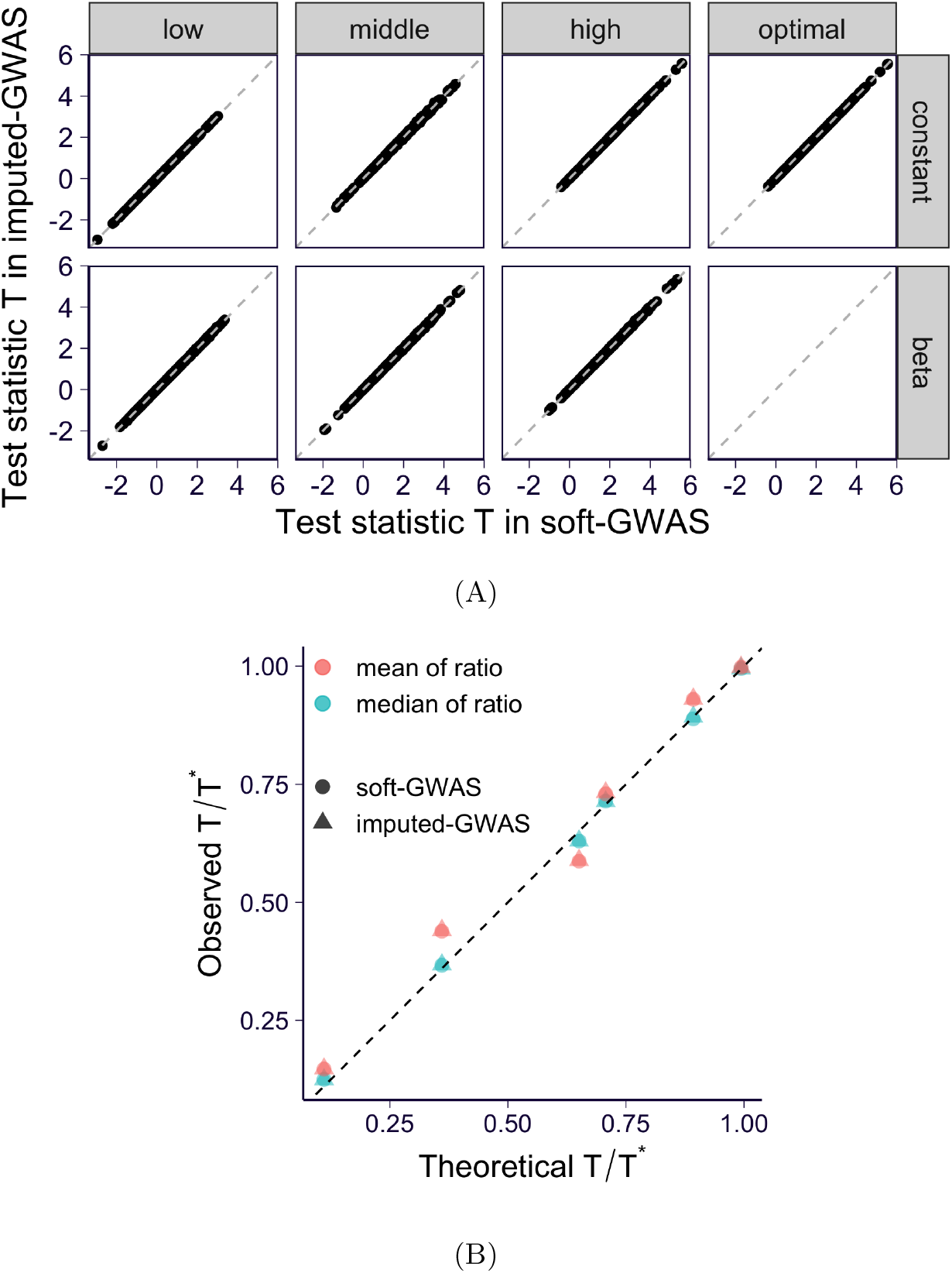
The results of the proposed GWAS approaches, imputed-GWAS and soft-GWAS, on the simulated data. **(A)** The test statistics are shown by panels where each panel presents the results under a specific *γ* distribution (the distribution type is organized in rows and the accuracy of *γ* is organized in columns). The test statistics of soft-GWAS are shown on x-axis and the ones of the imputed-GWAS are shown on y-axis. The gray dashed line is *y* = *x*. **(B)** The observed and theoretical relative power are shown. For each simulation setting, the theoretical result is calculated based on Eq 14. And the observed ratio is calculated for all replications separately and the median (in green) and mean (in red) across all replications are shown. Circle points indicate the results from soft-GWAS and the triangular ones indicate the results from imputed-GWAS. The black dashed line is *y* = *x*.

Furthermore, we examined the bias and power introduced by soft-GWAS and imputed-GWAS. Here we measured the relative power by the ratio of test statistic over the test statistic in “optimal” case, *T*\*. We found that the observed biases nicely agree with the theoretical result described in Section 2.2 (Supplementary Figure 5a). Moreover, And similarly, the relative powers also agree with the theoretical result (Figure 3B and Supplementary Figure 5b). So, in practice, before running the GWAS, we can get a good insight about the potential bias and power gain upper bound via these theoretical results.

### 3.3 Applying the imputation scheme to trios Framingham transcriptomic study

To investigate the utility of the proposed haplotype imputation scheme, we ran the scheme on the genotype and gene expression profile from Framingham Heart Study (Section 2.5). We selected Framingham data for four reasons: 1) it has the genotype data for the trio so we know the actual haplotype origin as the gold standard; 2) it has the whole transcriptome data so we observe hundreds of parental phenotypes for each chromosome; 3) the cis-regulation of gene expression captures a substantial amount of heritability so that we can rely on a local genetic model to avoid multiple-chromosome fitting; 4) it is straightforward to apply PRS-based imputation to gene expression since there are many high-quality and publicly available transcriptome predictors.

Overall, we extracted 266 trios and applied the imputation scheme to the child’s haplotype and parental transcriptome. As we phased the genotype data using the trio information so the first haplotype of the child is, by design, from father. To verify this result, we calculated the genetic relatedness between the child’s haplotypes and the parents’ genotypes. The observed relatedness is consistent with the expectation (Section 2.5 and Supplementary Figure 6). So, *γ* = 1 for all individuals is the ground truth of the imputation.

As for the PRS-based approach, we used publicly available gene expression models as described in Section 2.5. For each chromosome, there are about 80-950 genes per chromosomes available for imputation. The PRS-based approach using EN models achieves good accuracy in all chromosomes and the quality is higher for those chromosomes that have more genes (Figure 4). For many individuals, the imputation output *γ* is very close to 1 which is the ground truth with some strong errors which has *γ* very close to 0. And the same trend is observed in the imputation runs using EN DAPG models as well (Supplementary Figure 7) As we downsampled the genes for each chromosome, the imputation becomes less certain in the sense that the *γ* values are pushed towards the middle (Supplementary Figure 8 and 9). Since the imputation quality affects the downstream imputed-GWAS analysis, we calculated the expected power relative to no imputation scenario (*γ* = 0.5) which corresponds to the GWAX approach (Figure 5). As expected, the power increases when more genes are used. Moreover, the imputation using EN models performs better than the one using EN DAPG models. It could be attributed to the fact that EN DAPG is a sparse model so EN DAPG models are more sensitive to the imperfect variant overlapping between the HRC v1.1 and GTEx v8 (which is the variant panel used in EN and EN DAPG models). Roughly speaking, the imputed-GWAS is more powerful than the GWAX approach when the number of genes used in the imputation exceeds 100. In other word, we need more than 100 parental phenotypes to obtain a good enough imputation such that imputed-GWAS outperforms GWAX. As the current UK Biobank has only 12 parental phenotypes, we conclude that we have too few parental phenotypes to support the current imputation scheme.

**Figure 4:**
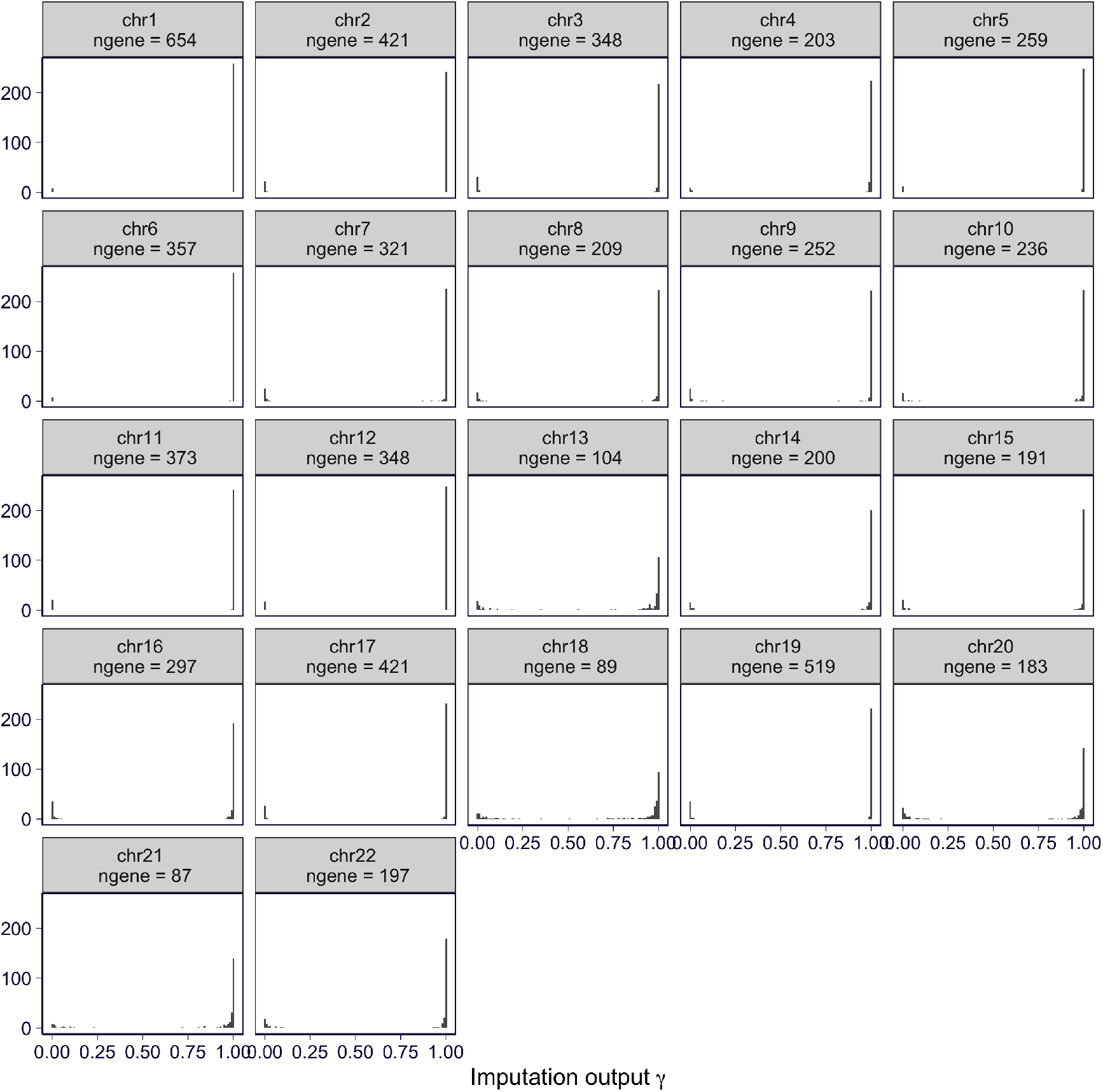
PRS-based imputation results using EN DAPG models as the genetic predictor. The histogram of the imputation output *γ* (ground truth is 1) is shown for each chromosome in the panels. “ngene” represents the number of genes used in the imputation.

**Figure 5:**
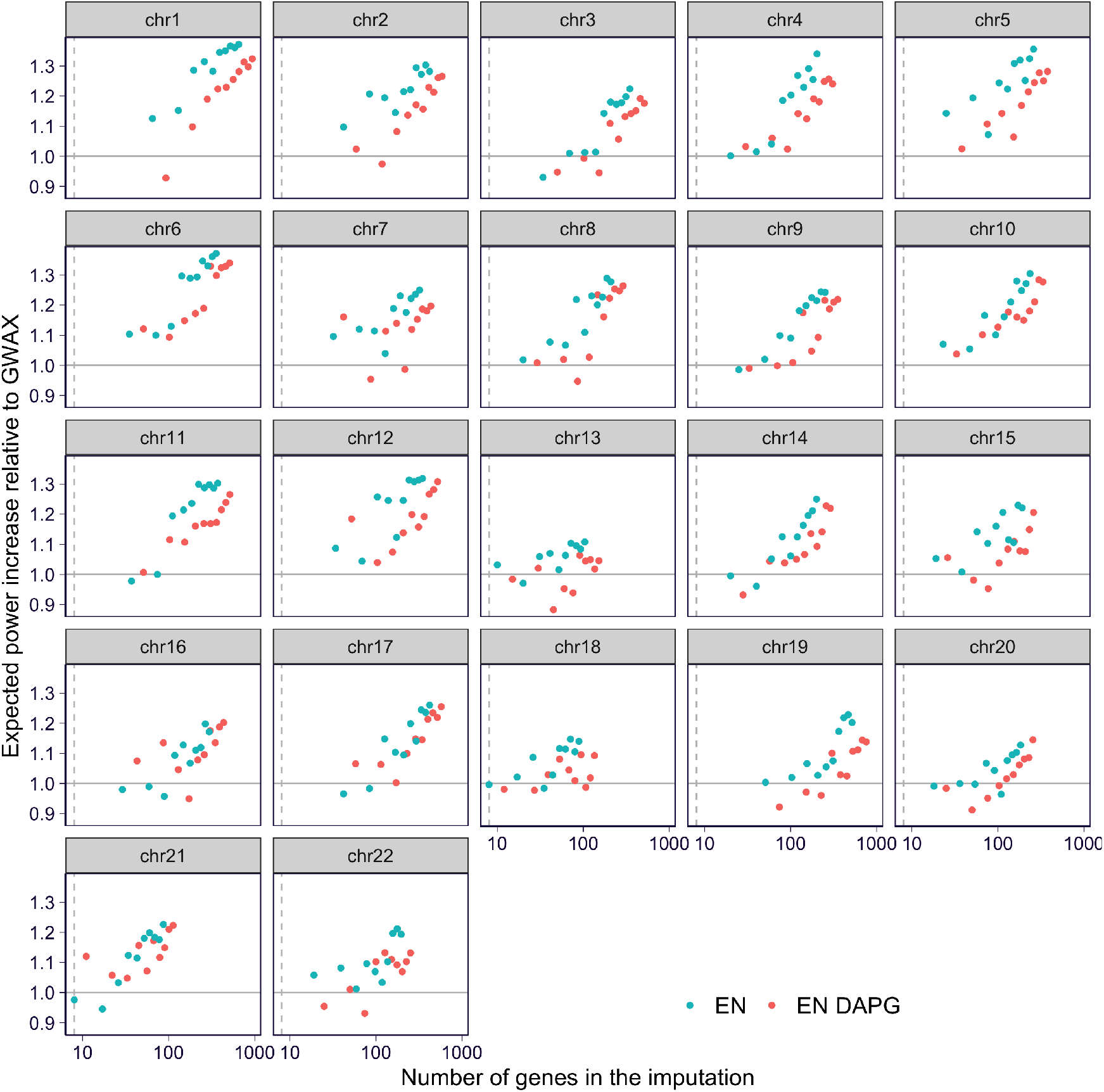
The expected power increase of the imputed-GWAS using the PRS-based imputation relative to the approach without imputation (GWAX). For each PRS-imputation run on either full data or downsampled data (downsampling genes), the number of genes used in the imputation is plotted on x-axis. And on y-axis, we show the corresponding relative power of the GWAS which uses the imputation result as input. In particular, the relative power is defined as the expected test statistic using the imputation over the expected test statistic using *γ* = 0.5 (*i.e*. the GWAX approach where no imputation is integrated). The results using EN models as genetic predictor are shown in green, and the ones using EN DAPG models are shown in red. The horizontal solid line is *y* = 1 corresponding to no power increase relative to GWAX. And the vertical dashed line is *x* = 10 which is approximately the number of parental phenotypes available in the UK Biobank.

## 4 Discussion

In this paper, we propose a two-approach to perform genotype-phenotype association study in the situation that we only have access to child’s genotype and parental phenotype. Specifically, in the first step, we propose a likelihood based imputation scheme to infer the parental origin of the child’s haplotypes where the statistical model focuses on the non-focal parental phenotypes. And, in the downstream, we propose an efficient approach to integrate the imputation results to association test, which we call imputed-GWAS.

In the simulation study, we show that the imputation scheme can capture the parental origin when we observe tens to hundreds of heritable parental phenotypes. Furthermore, we show, theoretically and via simulation, that the power of the imputed-GWAS relies on the quality of the imputation. When the imputation is noisy, the imputed-GWAS doesn’t typically benefit from such imputation.

Lastly, we perform the haplotype imputation on transcriptome and genotype data in trios from Framingham Heart Study. Here the imputation is performed within each chromosome and there are tens to hundreds of genes per chromosome. In this specific case, we show that the PRS-based imputation scheme is able to effectively impute the haplotype origin. And we observe that we need at least around 100 genes to have theoretical power gain in the downstream imputed-GWAS analysis (relative to the GWAS without no imputation, *e.g*. GWAX). From here, we conclude that this two-step approach is not applicable to the current biobank scale data set due to the lack of parental phenotypes.

Our paper has many limitations both in the proposed approach and in the data analysis. First, in the two-step procedure, we assume that all phenotypes are independent. This assumption simplifies the genetic model used in the imputation and the current framework is easily extendable to consider corrected non-focal phenotypes. However, the other complication comes from the correlation between the non-focal phenotypes and the focal phenotype (GWAS phenotype). It potentially introduces biases or false positives/negatives in the downstream GWAS, which we don’t study in details in the paper. Secondly, some of our conclusions are based on simulation where all the limitations about simulation study apply. Especially, in our case, our conclusion about power and performance is always dependent on parameter settings such as genetic architecture of the trait, heritability, and etc. Even though we simulate under carefully picked parameters so that it is not too far from the reality but someone should always interpret these results with cautious. Finally, due to the lack of data, we don’t have a complete run of the two-step procedure on real data. With this, we want to emphasize the importance of parental phenotypes and we urge the development of a more systematic procedure for parental phenotype collection. In the context of studying late-onset disease, a rich set of parental phenotypes is of great importance. And, along this line, our paper provides an example that how these parental phenotypes can facilitate the research and enable us to dig deeper into the data.

## 5 Code Availability

The code used for this work is at https://github.com/liangyy/haplotype-po.

## 6 Acknowledgement

We thank Hae Kyung Im for helpful discussion. Framingham data accession phs000007.v32.p1 was used. The Framingham Heart Study is conducted and supported by the National Heart, Lung, and Blood Institute (NHLBI) in collaboration with Boston University (Contract No. N01-HC-25195). This manuscript was not prepared in collaboration with investigators of the Framingham Heart Study and does not necessarily reflect the opinions or views of the Framingham Heart Study, Boston University, or NHLBI. Funding for SHARe Affymetrix genotyping was provided by NHLBI Contract N02-HL64278. SHARe Illumina genotyping was provided under an agreement between Illumina and Boston University. Additional funding for SABRe was provided by Division of Intramural Research, NHLBI, and Center for Population Studies, NHLBI. This research has been conducted using the UK Biobank Resource under Application Number 19526.

## Notes

### Competing Interest Statement

The authors have declared no competing interest.

### Summary of Updates

The acknowledgment has been updated.

