## Supplementary Figures and Notes for "Imputing the parental origin of the sibling’s haplotype from parental phenotypes"

### Supplementary Figures and Notes: Imputing the parental origin of the sibling's haplotype from parental phenotypes

Yanyu Liang[1,*]

**1** Section of Genetic Medicine, University of Chicago, Chicago, Illinois, United States of America

9 PRS-based imputation results on the downsampled data using EN DAPG models as the genetic predictor. 10

### 1 Supplementary Figures

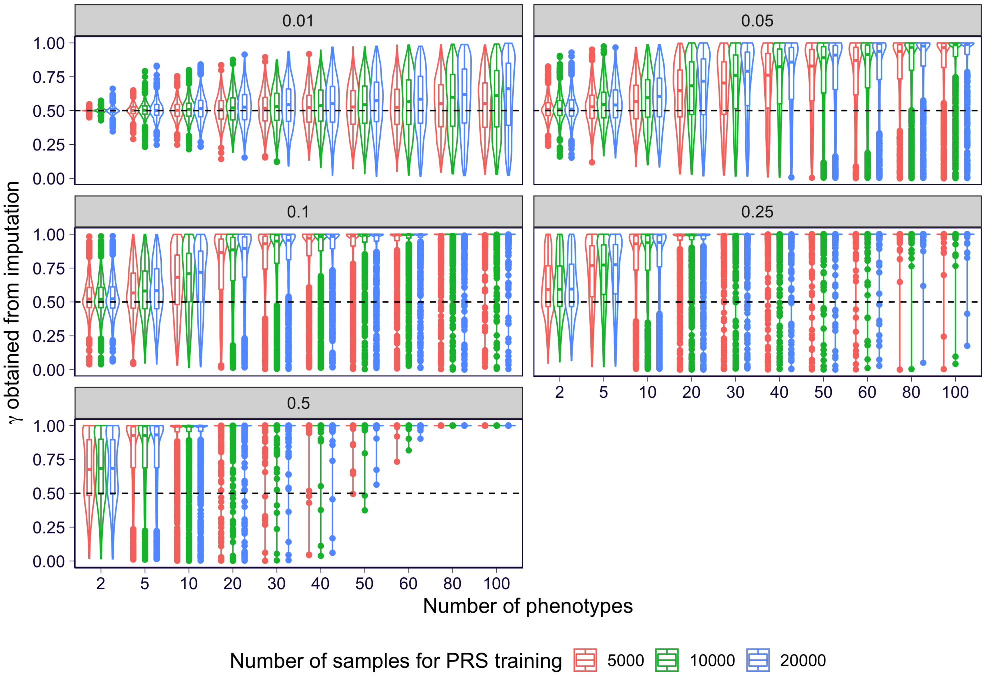

**Supplementary Figure 1. The imputation performance of the PRS-based approach with PRS trained with different sample sizes.** The imputation performance under a different heritability is shown in each panel. Within each panel, the number of phenotypes included in the imputation is shown on x-axis and the imputation accuracy (the probability that the posterior $Z$ is correctly assigned) is shown on the y-axis. The violin/boxplot contains the results on all of the 1,000 individuals included in the imputation. The results of the PRS-based approaches that are based on PRSs trained in 5,000, 10,000, and 20,000 individuals are colored in red, blue, and green respectively.

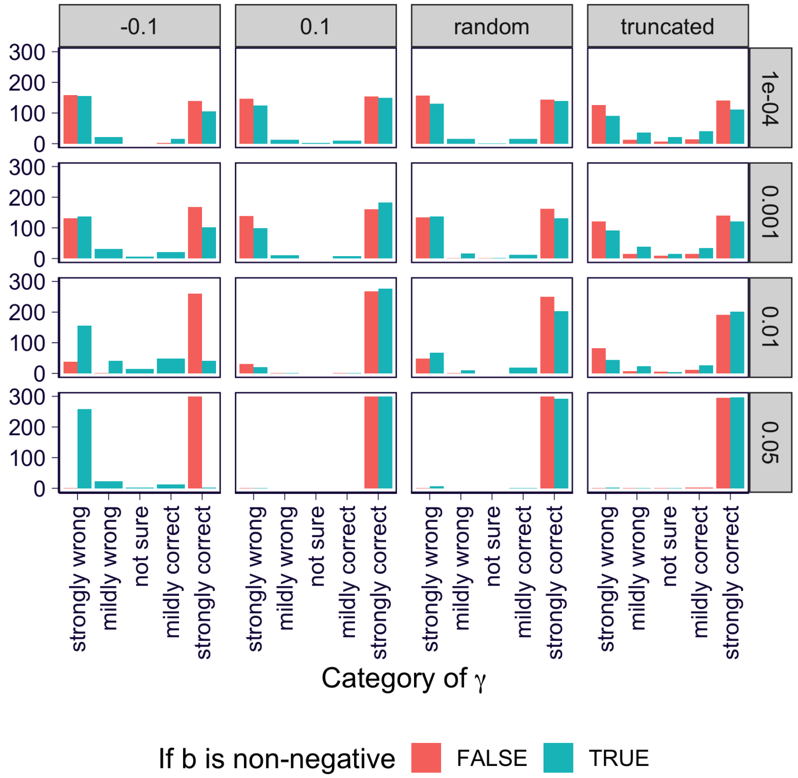

**Supplementary Figure 2. Comparing the performance of PRS-based approach with/without non-negative constraint on the coefficient.** The simulated data used here is generated in the context of using parental transcriptome (second scheme in Section 2.3) with sample size = 300 and number of genes = 500. Each panel shows the imputation results under one simulation setting with per-gene heritability organized in rows and the distribution of the true $b$ in columns. Specifically, the panels labeled with "random" mean that $b \sim N(0, 1)$ so that it has random sign and, similarly, the ones with "truncated" mean that $b \sim \max(0, N(0, 1))$ so that it is non-negative. For illustration purpose, we binned the imputation results into 5 categories on x-axis, in which $r$ (the probability of being correct) are binned into $[0, 0.1)$, $[0.1, 0.4)$, $[0.4, 0.6]$, $(0.6, 0.9]$, and $(0.9, 1]$ representing "strongly wrong", "mildly wrong", "not sure", "mildly correct", and "strongly correct" respectively. And y-axis shows the count of the bin. The results of the PRS-based approaches with and without non-negative constraint on $b$ are shown in green and red.

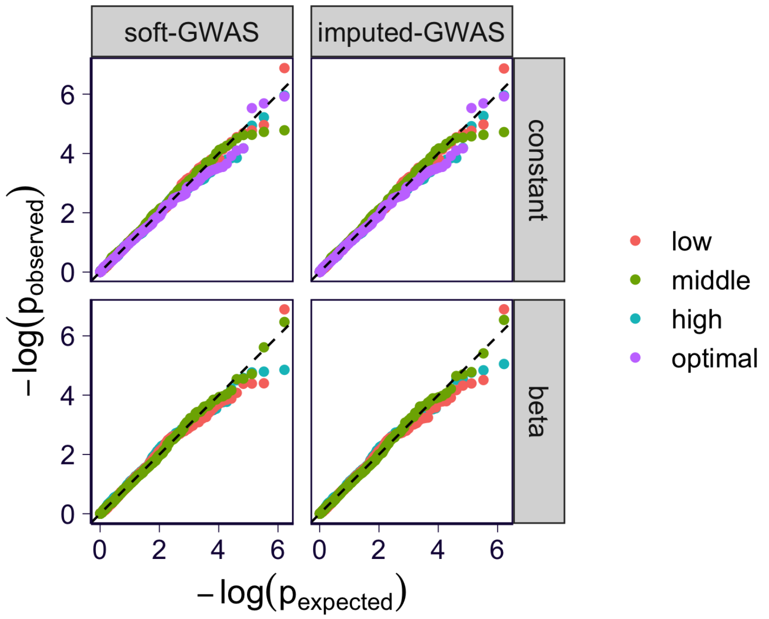

**Supplementary Figure 3. QQ-plot of the proposed GWAS tests under the simulated null data.** QQ-plot of the association p-values (expected on axis versus observe on y-axis in $-\log$ scale) are shown. Each panel shows the results on one GWAS approach (soft-GWAS or imputed-GWAS in columns) and one distribution of $\gamma$ (constant or beta in rows). The QQ-plots corresponding to different $\gamma$ distributions are drawn and shown separately in different colors (see more details at Section 2.4).

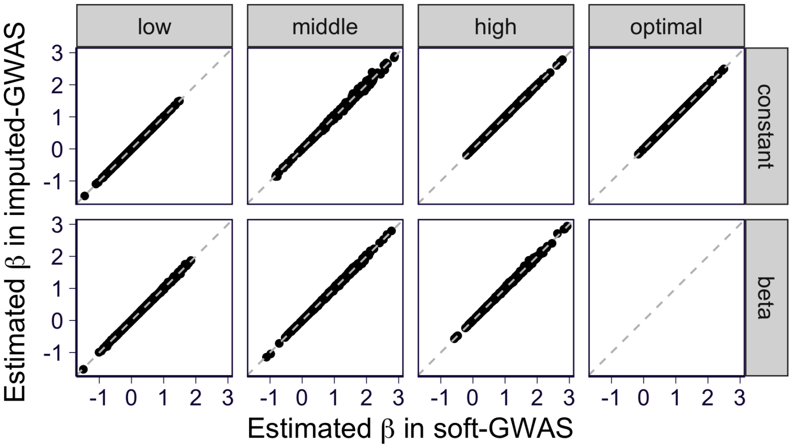

**(a)**

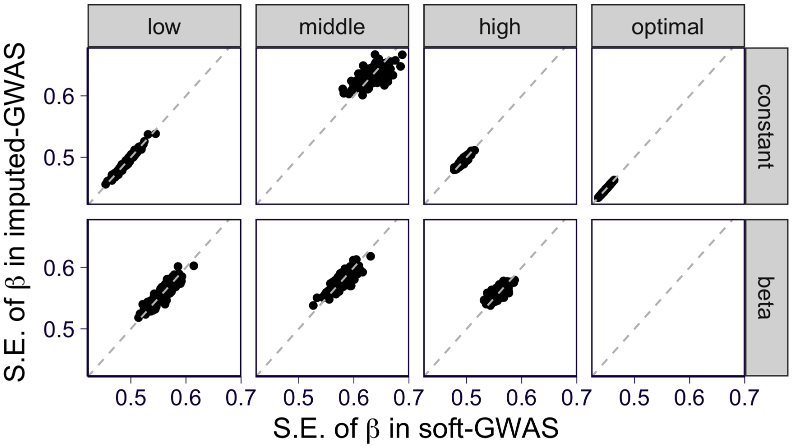

**(b)**

**Supplementary Figure 4. Comparing the effect size estimates in soft-GWAS and imputed-GWAS on simulated data.** Each panel presents the results under a specific $\gamma$ distribution (the distribution type is organized in rows and the accuracy of $\gamma$ is organized in columns). The results of soft-GWAS are shown on x-axis and the ones of the imputed-GWAS are shown on y-axis. The gray dashed line is $y = x$. **(A)** The estimated effect sizes $\hat{\beta}$ are shown. **(B)** The standard error of $\hat{\beta}$ are shown.

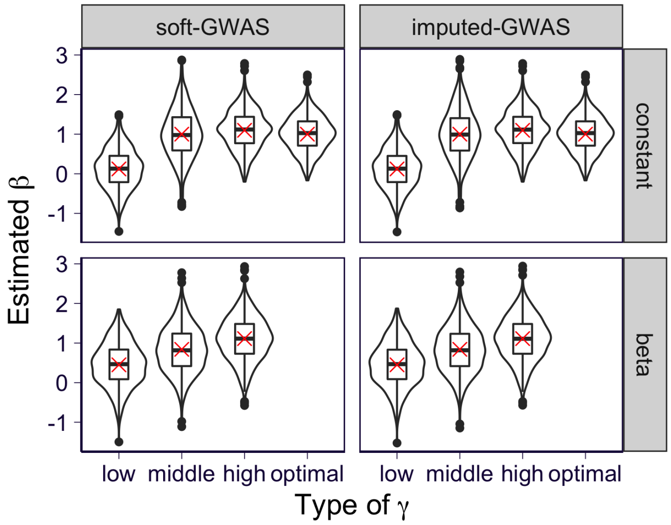

**(a)**

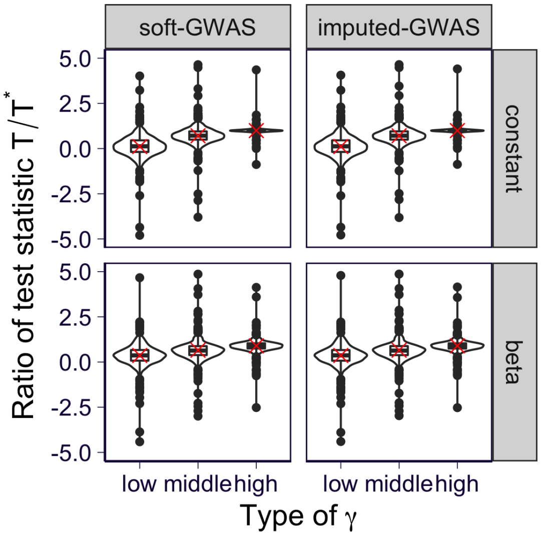

**(b)**

**Supplementary Figure 5. Comparing the theoretical and observed bias and relative power.** Each panel presents the results under a specific $\gamma$ distribution (the distribution type is organized in rows and the type of method is organized in columns). And the results are stratified by the accuracy of $\gamma$ on x-axis. **(A)** The estimated effect sizes $\hat{\beta}$ are shown in the violin/boxplot and the red cross indicates the expected effect size after accounting for the theoretical bias. **(B)** The observed across all replications are shown in the violin/boxplot where outliers with observed ratio outside $[-5, 5]$ are excluded. The red cross shows the theoretical relative power.

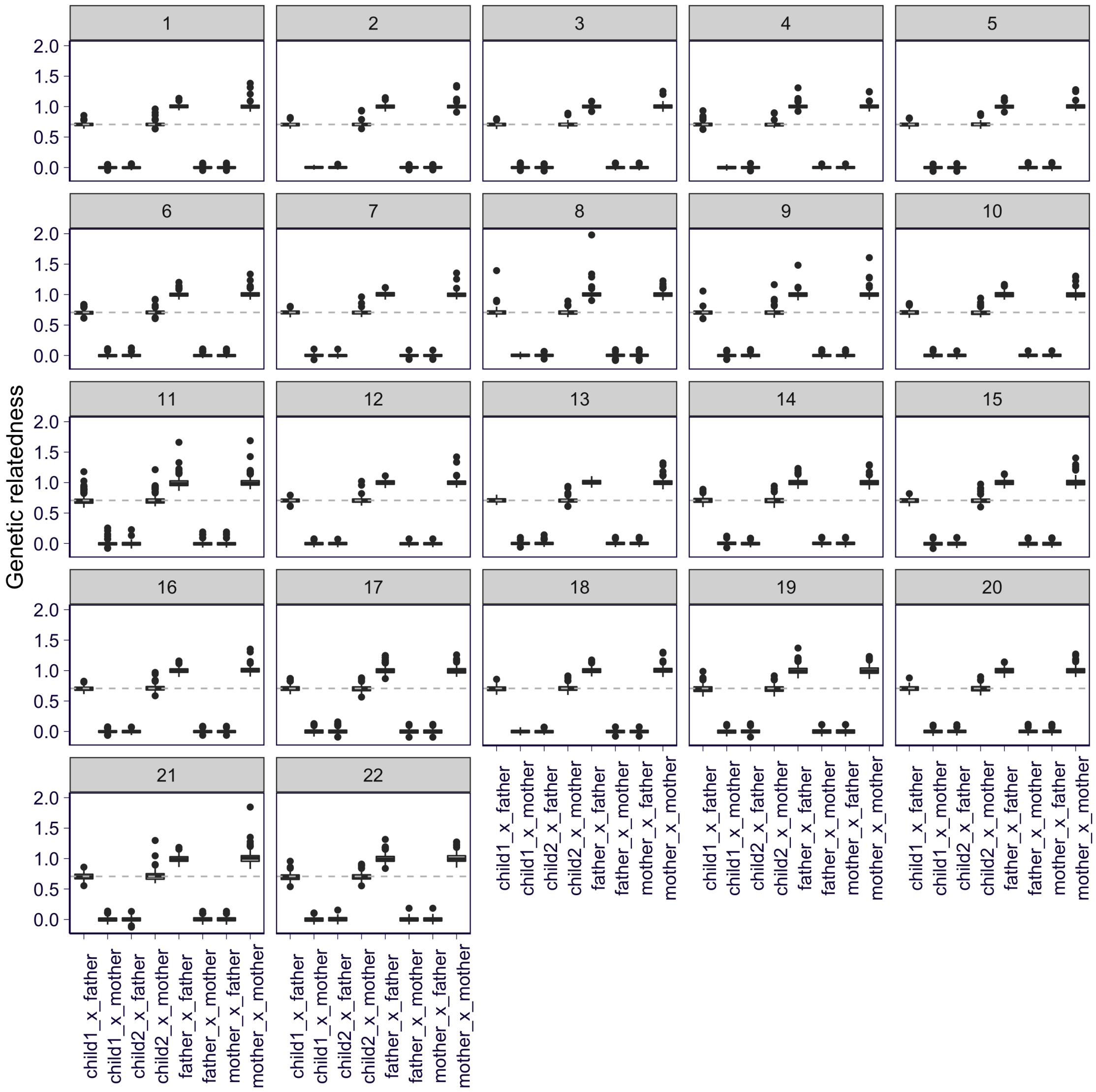

**Supplementary Figure 6. Genetic relatedness between child's haplotypes and parents' genotypes in Framingham trios.** Each panel presents the results on one of the 22 autosomes. On x-axis, "sample1_x_sample2" means the genetic relatedness between sample1 and sample2 where sample1 and sample2 can be either haplotype or genotype. "child1" represents the first haplotype of the child and "child2" means the second one. "father" and "mother" represent the genotype of the father and mother. On y-axis, the genetic relatedness is shown (see detailed definition in Section 2.5). The horizontal dashed line is $y = \frac{1}{\sqrt{2}}$ which is the expected genetic relatedness between a haploid and a diploid if the haploid comes from the diploid.

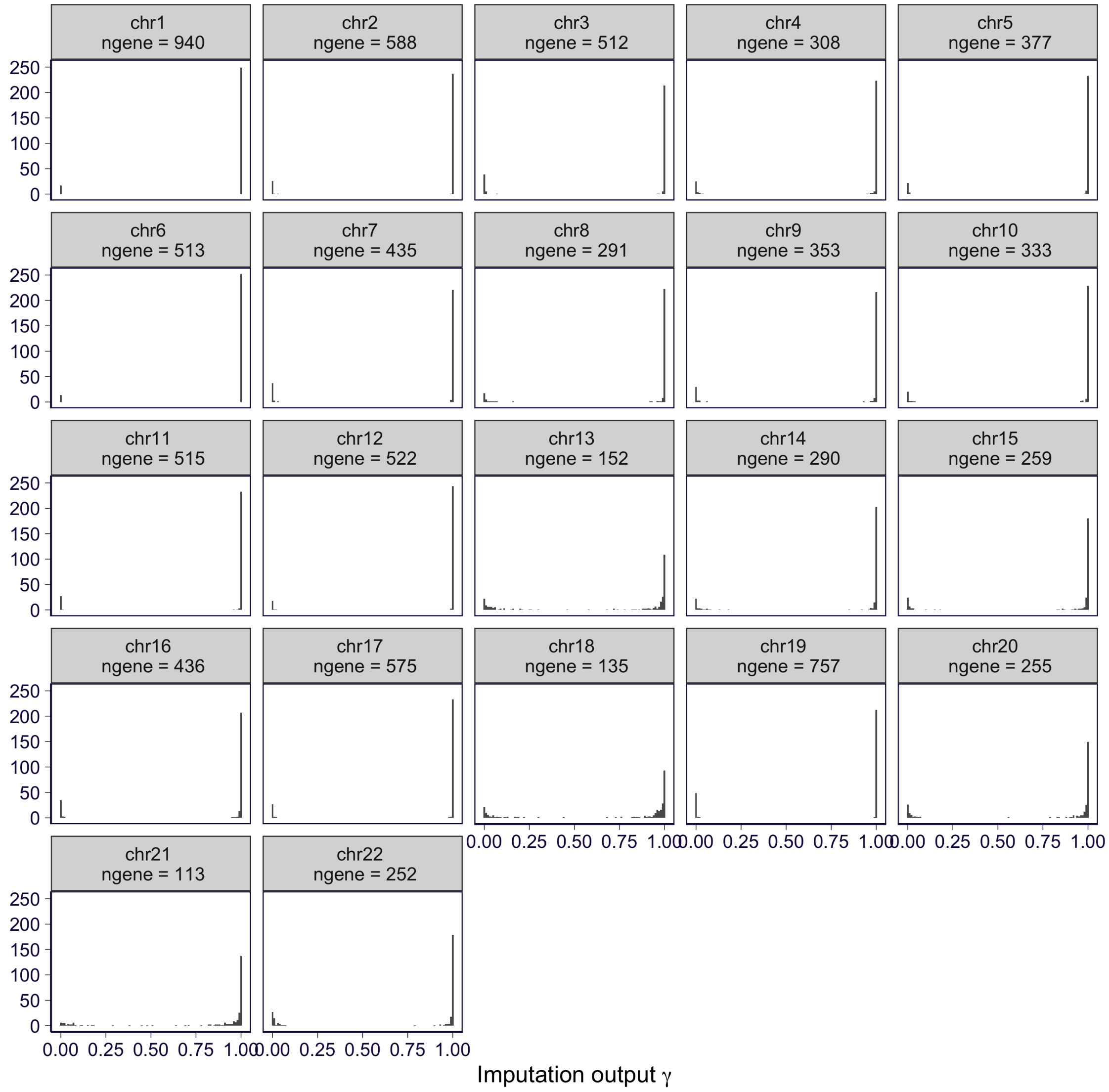

**Supplementary Figure 7. PRS-based imputation results using EN DAPG models as the genetic predictor.** The histogram of the imputation output $\gamma$ (ground truth is 1) is shown for each chromosome in the panels. "ngene" represents the number of genes used in the imputation.

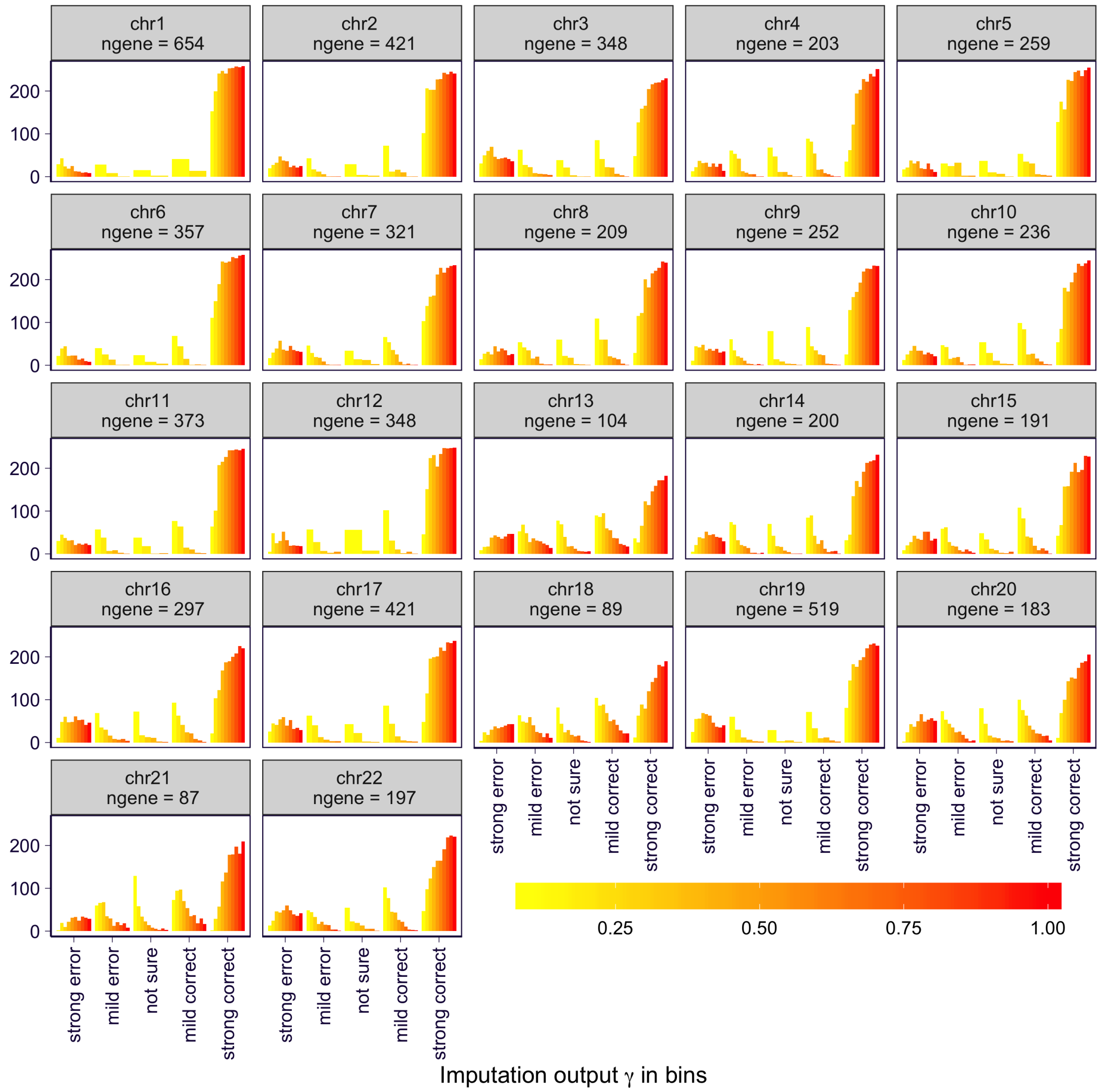

**Supplementary Figure 8. PRS-based imputation results on the downsampled data using EN models as the genetic predictor.** The imputation results on the downsampled data is shown for each chromosome in the panels. The imputation output $\gamma$ is stratified into 5 bins $[0, 0.1)$, $[0.1, 0.4)$, $[0.4, 0.6]$, $(0.6, 0.9]$, and $(0.9, 1]$ representing "strongly wrong", "mildly wrong", "not sure", "mildly correct", and "strongly correct" respectively. And y-axis shows the count of the bin. The results are colored by the downsampling fraction relative to the full data. "ngene" represents the number of genes in the full data.

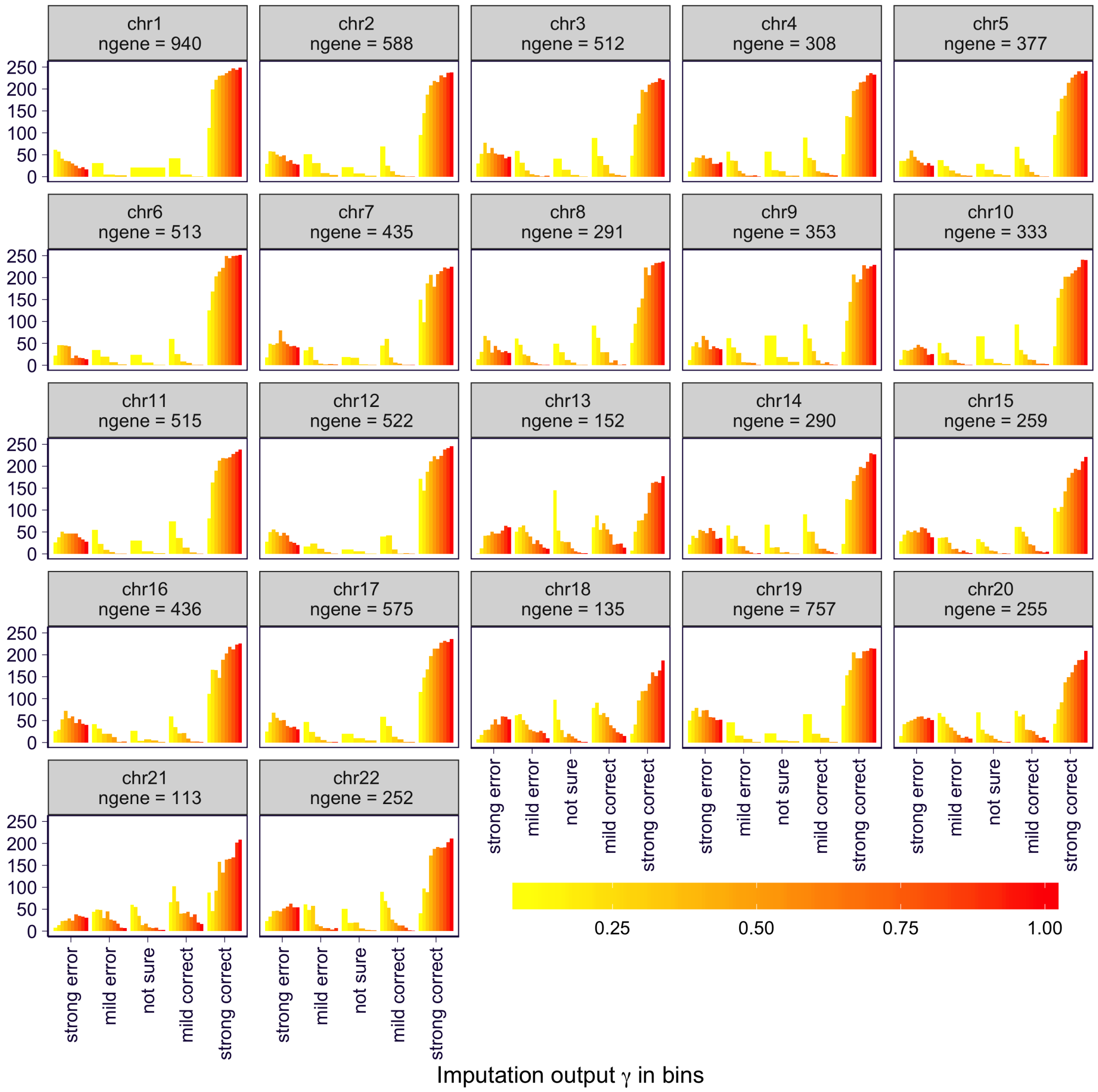

**Supplementary Figure 9. PRS-based imputation results on the downsampled data using EN DAPG models as the genetic predictor.** The imputation results on the downsampled data is shown for each chromosome in the panels. The imputation output $\gamma$ is stratified into 5 bins $[0, 0.1)$, $[0.1, 0.4)$, $[0.4, 0.6]$, $(0.6, 0.9]$, and $(0.9, 1]$ representing "strongly wrong", "mildly wrong", "not sure", "mildly correct", and "strongly correct" respectively. And y-axis shows the count of the bin. The results are colored by the downsampling fraction relative to the full data. "ngene" represents the number of genes in the full data.

#### 2 Supplementary Notes

##### 2.1 The EM algorithm to impute haplotype origin

Based on Eq 9, we have ($p$ indexes phenotypes and $i$ indexes individuals)

$$
\begin{aligned}
Q(\theta, \theta^{(t)}) &:= \mathrm{E}_{Z|y,H,\theta^{(t)}}[\log \Pr(y^{\text{father}}, y^{\text{mother}}, Z|H^1, H^2; \theta)] && (1)\\
&= \sum_i \mathrm{E}_{Z_i|y_i,H_i,\theta^{(t)}}[\log \Pr(Z_i) + \sum_p \log \Pr(y_{i,p}^{\text{father}}, y_{i,p}^{\text{mother}}|Z_i, H_i^1, H_i^2; \theta)] && (2)\\
&\text{(assume phenotypes are independent conditioning on haplotypes)} && (3)\\
&= const. + \sum_i \mathrm{E}_{Z_i|y_i,H_i,\theta^{(t)}}[\sum_p \log \Pr(y_{i,p}^{\text{father}}, y_{i,p}^{\text{mother}}|Z_i, H_i^1, H_i^2; \theta)] && (4)\\
&\text{(as prior knowledge, we have } \Pr(Z_i) = 0.5\text{)} && (5)
\end{aligned}
$$

Since $Z$ indicates if haplotype 1 is from father, we have

$$
\begin{aligned}
&\log \Pr(y_{i,p}^{\text{father}}, y_{i,p}^{\text{mother}}|Z_i, H_i^1, H_i^2; \theta) && (6)\\
=&Z_i F_1(y_{i,p}^{\text{father}}, y_{i,p}^{\text{mother}}, H_i^1, H_i^2) + (1 - Z_i) F_2(y_{i,p}^{\text{father}}, y_{i,p}^{\text{mother}}, H_i^1, H_i^2) && (7)
\end{aligned}
$$

where

$$
\begin{aligned}
F_1(y_{i,p}^{\text{father}}, y_{i,p}^{\text{mother}}, H_i^1, H_i^2) &:= \log \Pr(y_{i,p}^{\text{father}}, y_{i,p}^{\text{mother}}|Z_i = 1, H_i^1, H_i^2) && (8)\\
&= l(y_{i,p}^{\text{father}}, H_i^1) + l(y_{i,p}^{\text{mother}}, H_i^2) && (9)\\
F_2(y_{i,p}^{\text{father}}, y_{i,p}^{\text{mother}}, H_i^1, H_i^2) &:= \log \Pr(y_{i,p}^{\text{father}}, y_{i,p}^{\text{mother}}|Z_i = 0, H_i^1, H_i^2) && (10)\\
&= l(y_{i,p}^{\text{father}}, H_i^2) + l(y_{i,p}^{\text{mother}}, H_i^1) && (11)
\end{aligned}
$$

and $l(y, H)$ represent the log-likelihood of observing phenotype $y$ and haplotype $H$. For simplicity, we use $F_1(i,p)$ and $F_2(i,p)$ as the short form of $F_1(y_{i,p}^{\text{father}}, y_{i,p}^{\text{mother}}, H_i^1, H_i^2)$ and $F_2(y_{i,p}^{\text{father}}, y_{i,p}^{\text{mother}}, H_i^1, H_i^2)$.

Combining Eq 4 and 7, we have

$$
\begin{aligned}
Q(\theta, \theta^{(t)}) &= const. + \sum_i \mathrm{E}_{Z_i|y_i,H_i,\theta^{(t)}}[\sum_p Z_i F_1(i,p) + (1 - Z_i) F_2(i,p)] && (12)\\
&= const. + \sum_i w_i \sum_p F_1(i,p) + \sum_i (1 - w_i) \sum_p F_2(i,p) && (13)
\end{aligned}
$$

where $w_i := \mathrm{E}_{Z_i|y_i,H_i,\theta^{(t)}}[Z_i] = \Pr(Z_i = 1|y_i, H_i, \theta^{(t)})$.

At E step, we update $w_i$ by applying Bayes rule

$$
\begin{aligned}
w_i &= \frac{\Pr(y_i|Z_i = 1, H_i, \theta^{(t)}) \Pr(Z_i = 1)}{\sum_{k \in \{0,1\}} \Pr(y_i|Z_i = k, H_i, \theta^{(t)}) \Pr(Z_i = k)} && (14)\\
&= \frac{\exp(\sum_p F_1(i,p;\theta^{(t)}))}{\exp(\sum_p F_1(i,p;\theta^{(t)})) + \exp(\sum_p F_2(i,p;\theta^{(t)}))} && (15)
\end{aligned}
$$

where, notice that, we need to use $\theta^{(t)}$ when evaluating $F_1$ and $F_2$.

At M step, we update $\theta$ by

$$\theta = \arg\max_\theta \sum_i w_i \sum_p F_1(i, p; \theta) + \sum_i (1 - w_i) \sum_p F_2(i, p; \theta) \tag{16}$$

Specifically, for on-the-fly approach (Eq 5), it corresponds to solving weighted least squares. For instance, to obtain father-specific model parameters $\beta^{\text{father}}$ and $\sigma^2_{\text{father}}$, it is equivalent to solve weighted least squares with the following settings

$$\text{response} = \begin{bmatrix} y^{\text{father}} \\ y^{\text{father}} \end{bmatrix} \tag{17}$$

$$\text{design matrix} = \begin{bmatrix} H^1 \\ H^2 \end{bmatrix} \tag{18}$$

$$\text{weight} = \begin{bmatrix} w \\ 1 - w \end{bmatrix} \tag{19}$$

Similarly, to obtain mother-specific model parameters $\beta^{\text{mother}}$ and $\sigma^2_{\text{mother}}$, we solve with

$$\text{response} = \begin{bmatrix} y^{\text{mother}} \\ y^{\text{mother}} \end{bmatrix} \tag{20}$$

$$\text{design matrix} = \begin{bmatrix} H^1 \\ H^2 \end{bmatrix} \tag{21}$$

$$\text{weight} = \begin{bmatrix} 1 - w \\ w \end{bmatrix} \tag{22}$$

For the PRS based approach (Eq 6), we can simply replace $H_1$ and $H_2$ with the haplotypic PRS $\widetilde{G}^1$ and $\widetilde{G}^2$ and the similar calculation can be applied to solve for father/mother-specific $b^p$ and other parameters. Since we assume $b^p$ to be non-negative, at the M step, we set $b^p = 0$ if the weight least squares gives negative $b^p$.

#### 2.2 Fitting multiple chromosomes in iterative manner

The haplotype phasing is done for each chromosome. So, the EM algorithm in Section 2.1 can only handle one chromosome since the likelihood requires phased haplotypes. Even though one could fit one chromosome at a time while treating the contribution from other chromosomes as noise, to fit all chromosomes jointly is preferred. The reason is that the former captures at most per-chromosome heritability while the latter models the contribution from all chromosomes which, in other words, reduces the noise.

In principle, to handle multi-chromosome situation, instead of having $Z_i$ being a binary value, $Z_i$ should be a vector with each entry representing the parental origin of the corresponding chromosome. This creates complexities in the E step of the EM algorithm. At the E step, one needs to sum over all the possible configurations of $Z_i$. The number of terms in this summation is $2^2 2 = 4,194,304$, which is tractable but introduces heavy computation.

To avoid such computational burden, we apply an alternative approach which considers the contributions from other chromosomes while still handling one chromosome at a time. In this approach, we loop over chromosomes. At each chromosome, we apply the EM algorithm and then we update $y^{\text{father}}$ and $y^{\text{mother}}$ to the residual of the current model. In other words, the procedure is:

1. Initialization: $\beta = 0$ (or $b = 0$) and $Z^1 = \cdots = Z^{22} = 0.5$.

2. For chromosome 1 to 22, do the following until convergence

   (a) Run EM updates until convergence and update $Z^k$ correspondingly.

   (b) Let $y^{\text{father}} \leftarrow y^{\text{father}} - \hat{y}^{\text{father}}$ and $y^{\text{mother}} \leftarrow y^{\text{mother}} - \hat{y}^{\text{mother}}$ where $\hat{y}$ is the predicted values of the current EM fit.

3. Return $Z^1, \cdots, Z^{22}$

#### 2.3 The algorithm for soft-GWAS

In this paper, we are interested in testing the association between a focal phenotype and each of the SNPs. The likelihood of the GWAS problem is described in Eq 10 where $H^1$ and $H^2$ represent the haplotypes of a single SNP. And from the imputation results, we have $\Pr(Z_i = 1) = \gamma_i$. Let's further assume that $y^\pi = H^\pi \beta + e,\ e \sim N(0, \sigma^2)$. The model parameters can be estimated by maximum likelihood estimation

$$\hat{\beta}, \hat{\sigma}^2 = \arg\max_{\beta, \sigma^2} \Pr(y^\pi | H^1, H^2; \beta, \sigma^2) \tag{23}$$

And we can construct a likelihood ratio test as follow

$$\lambda_{\text{LR}} = -2[\log \Pr(y^\pi | H^1, H^2; \beta = 0, \sigma^2 = \hat{\sigma}_0^2) - \log \Pr(y^\pi | H^1, H^2; \beta = \hat{\beta}, \sigma^2 = \hat{\sigma}^2)] \tag{24}$$

where $\hat{\sigma}_0^2 := \arg\max_{\sigma^2} \Pr(y^\pi | H^1, H^2; \beta = 0, \sigma^2)$.

Notice that solving Eq 23 is similar to the on-the-fly approach as described in Section 2.1. Here we need to replace the full-chromosome haplotypes in the on-the-fly approach with single-SNP haplotype and use $\Pr(Z_i) = \gamma_i$ instead of $\Pr(Z_i = 1) = 0.5$. Furthermore, the current EM scheme can be adapted to handling binary trait with minor modifications.

#### 2.4 Derivation of the power and bias in imputed-GWAS

Without loss of generality, we assume that the imputed-GWAS is on $y^{\text{father}}\ \widetilde{X}$ where $\widetilde{X} = \gamma H_1 + (1-\gamma) H_2$ (here we use $H_k$ in place of $H^k$ to avoid potential ambiguity). And the true model (allelic test model) is $y^{\text{father}} = H_1 \beta + \epsilon$. So, the imputed-GWAS estimate is

$$\hat{\beta} := (\widetilde{X}' \widetilde{X})^{-1} (\widetilde{X}' y^{\text{father}}) \tag{25}$$

In the following, we derive $\mathrm{E}(\hat{\beta})$ and $\mathrm{Var}(\hat{\beta})$ in order to analyze the bias and power of the imputed-GWAS estimate.

$$\widetilde{X}'\widetilde{X} = (\Gamma H_1 + (I - \Gamma) H_2)' (\Gamma H_1 + (I - \Gamma) H_2) \tag{26}$$

$$\approx \sum_i \gamma_i^2 H_{1,i}^2 + \sum_i (1 - \gamma_i)^2 H_{2,i}^2 \tag{27}$$

$$(\text{since } H_1 \perp\!\!\!\perp H_2) \tag{28}$$

$$\approx H^2 [\ \sum_i \gamma_i^2 + (1 - \gamma_i)^2\ ] \tag{29}$$

$$(H \perp\!\!\!\perp \gamma \text{ and } H_1, H_2 \sim iid) \tag{30}$$
$$= H^2 N\overline{S} \tag{31}$$
$$(\text{let } S = \gamma^2 + (1-\gamma)^2) \tag{32}$$

where $\Gamma = \text{diag}(\gamma)$ and $N$ is the sample size of the imputed GWAS.

$$\widetilde{X}' y^{\text{father}} = [\ \Gamma H_1 + (I-\Gamma) H_2\ ]'(\ H_1\beta + \epsilon\ ) \tag{33}$$
$$= (\ H_1'\Gamma H_1 \beta + H_1'\Gamma\epsilon\ ) + (\ H_2'(I-\Gamma)H_1\beta + H_2'(I-\Gamma)\epsilon\ ) \tag{34}$$
$$\approx H_1'\Gamma H_1\beta + H_1'\Gamma\epsilon + H_2'(I-\Gamma)\epsilon \tag{35}$$
$$(\text{since } H_1 \perp\!\!\!\perp H_2) \tag{36}$$
$$\approx H^2 N\overline{\gamma}\beta + \widetilde{X}'\epsilon \tag{37}$$

Based on Eq 31 and 37, we have

$$\text{E}(\hat{\beta}) \approx \text{E}[\frac{H^2 N\overline{\gamma}\beta + \widetilde{X}'\epsilon}{H^2 N\overline{S}}] \tag{38}$$
$$= \frac{\overline{\gamma}}{\overline{S}}\beta \tag{39}$$
$$(\text{since } H_1 \perp\!\!\!\perp \epsilon \text{ and } H_2 \perp\!\!\!\perp \epsilon \text{ so that } \text{E}(\widetilde{X}'\epsilon) = 0) \tag{40}$$

$$\text{Var}(X'\epsilon) = \text{Var}(H_1'\Gamma\epsilon) + \text{Var}(H_2'(I-\Gamma)\epsilon) \tag{41}$$
$$(\text{since } H_1 \perp\!\!\!\perp H_2) \tag{42}$$
$$= \sum_i \gamma_i H_{1,i}^2 \text{Var}(\epsilon) + \sum_i (1-\gamma_i)^2 H_{2,i}^2 \text{Var}(\epsilon) \tag{43}$$
$$\approx H^2 \text{Var}(\epsilon)[\ \sum_i \gamma_i^2 + (1-\gamma_i)^2\ ] \tag{44}$$
$$(\text{since } H \perp\!\!\!\perp \gamma \text{ and } H_1, H_2 \sim iid) \tag{45}$$
$$= H^2 N\overline{S}\text{Var}(\epsilon) \tag{46}$$

$$\text{Var}(\hat{\beta}) = \text{Var}(\ \frac{\widetilde{X}' y^{\text{father}}}{\widetilde{X}'\widetilde{X}}\ ) \tag{47}$$
$$\approx \frac{1}{[\ H^2 N\overline{S}\ ]^2}\text{Var}(\widetilde{X}' y^{\text{father}}) \tag{48}$$
$$= \frac{1}{[\ H^2 N\overline{S}\ ]^2}\text{Var}(H^2 N\overline{S} + \widetilde{X}'\epsilon) \tag{49}$$
$$= \frac{1}{[\ H^2 N\overline{S}\ ]^2}\text{Var}(\widetilde{X}'\epsilon) \tag{50}$$
$$= \frac{1}{[\ H^2 N\overline{S}\ ]^2} H^2 N\overline{S}\text{Var}(\epsilon) \tag{51}$$

$$= \frac{\text{Var}(\epsilon)}{H^2 N \overline{S}} \tag{52}$$

Based on Eq 39 and 52, we have

$$\text{E}(T) := E[\frac{\hat{\beta}}{\sqrt{\widehat{\text{Var}}(\hat{\beta})}}] \tag{53}$$

$$\approx \frac{E(\hat{\beta})}{\sqrt{\text{Var}(\hat{\beta})}} \tag{54}$$

$$= \frac{\overline{\gamma}}{\overline{S}} \frac{\sqrt{H^2 N \overline{S}}}{\sqrt{\text{Var}(\epsilon)}} \beta \tag{55}$$

$$= \frac{\overline{\gamma}}{\sqrt{\overline{S}}} \underbrace{\sqrt{\frac{NH^2}{\text{Var}(\epsilon)}} \beta}_{\text{E}(T^\star) \text{ from } y^{father} \sim H_1} \tag{56}$$

$$= \frac{\overline{\gamma}}{\sqrt{\overline{S}}} E(T^\star) \tag{57}$$

Eq 57 follows since in the ideal case, $\widetilde{X} = H_1$, $\overline{S} = \overline{\gamma} = 1$ so that $\text{E}(T^\star) = \sqrt{\frac{NH^2}{\text{Var}(\epsilon)}} \beta$.
